# APOBEC3B-dependent kataegis and TREX1-driven chromothripsis in telomere crisis

**DOI:** 10.1101/725366

**Authors:** John Maciejowski, Aikaterini Chatzipli, Alexandra Dananberg, Titia de Lange, Peter J. Campbell

## Abstract

Chromothripsis and kataegis are frequently observed in cancer and can arise from telomere crisis, a period of genome instability during tumorigenesis when depletion of the telomere reserve generates unstable dicentric chromosomes^1–5^. Here we report on the mechanism underlying chromothripsis and kataegis using an in vitro telomere crisis model. We show that the cytoplasmic exonuclease TREX1, which promotes the resolution of dicentric chromosomes^4^, plays a prominent role in chromothriptic fragmentation. In absence of TREX1, the genome alterations induced by telomere crisis primarily involve Breakage-Fusion-Bridge cycles and simple genome rearrangements rather than chromothripsis. Furthermore, we show that the kataegis observed at chromothriptic breakpoints is the consequence of cytosine deamination by APOBEC3B. In addition, APOBEC3B increased the frequency of chromothriptic fragmentation, possibly due to strand breakage after cytosine deamination. These data reveal that chromothripsis and kataegis arise from a combination of nucleolytic processing by TREX1 and cytosine editing by APOBEC3B.

---

To model telomere crisis we used a previously established model system based on RPE-1 cells in which the Rb and p53 pathways are disabled with shRNAs and telomere fusions are generated with a doxycycline-inducible dominant negative allele of the TRF2 subunit of shelterin^4,6^. The dicentric chromosomes induced in this system persist through mitosis to form long (50-200 μm) DNA bridges that are generally resolved before the connected daughter cells enter the next S phase. Bridge resolution is accelerated by the exonucleolytic activity of TREX1, which accumulates on the DNA bridge after nuclear envelope rupture^4,7–9^ and results in formation of RPA-coated single-stranded (ss) DNA. Rearranged clonal cell lines isolated after progression through this in vitro telomere crisis showed frequent chromothripsis in a pattern similar to cancer: the chromothripsis events were limited to (parts of) chromosome arms rather than whole chromosomes^4,10^. Furthermore, as is the case for chromothripsis in cancer, the breakpoints showed kataegis with the hallmarks of APOBEC3 editing^4,11,12^. To determine whether TREX1 contributes to chromothripsis after telomere crisis, TREX1-deficient cell lines generated by CRISPR/Cas9 editing (hereafter TREX1 KOs) were subjected to telomere crisis alongside the TREX1-proficient T2p1 cell line and hundreds of clonal post-crisis descendants were isolated.

Since only a subset of the isolated clones are expected to have experienced telomere crisis^4^, initial identification of clones with genomic alterations was necessary. To determine whether low-pass whole genome sequencing (WGS) can identify relevant copy number changes evident at higher coverages, 17 post-crisis clones derived from T2p1 were analyzed at both 1x and 30x sequence coverage (Fig. 1a-d)^13,14^. Among chromosomes showing no copy number (CN) changes in 1x WGS analysis, 67% also did not show CN changes in high coverage WGS and 30% showed <4 CN changes (hereafter referred to as simple events) (Fig. 1b). Only 3% of chromosomes lacking evidence for CN changes in 1x WGS were found to contain ≥4 CN changes (hereafter referred to as complex events) in 30x WGS (Fig. 1b). The discrepancy in the segments missed by 1x WGS but reported in the 30x data is likely due to the conservative thresholds used for calling gains and losses in low coverage data. Of 37 chromosomes showing simple events in 1x WGS, 19 were found to contain complex events in 30x WGS (Fig. 1b). Overall, the 1x analysis had an acceptable false negative rate of <10% (32 of 391 chromosomes) with regard to identifying chromosomes with complex events. Similarly, the false positive rate of the 1x coverage analysis was well below 10%, since only 1 out of 19 chromosomes with complex events detected in 1x WGS did not show ≥4 CN in 30x coverage. These data indicated that 1x WGS allows identification of informative post-crisis clones.

**Figure 1.**
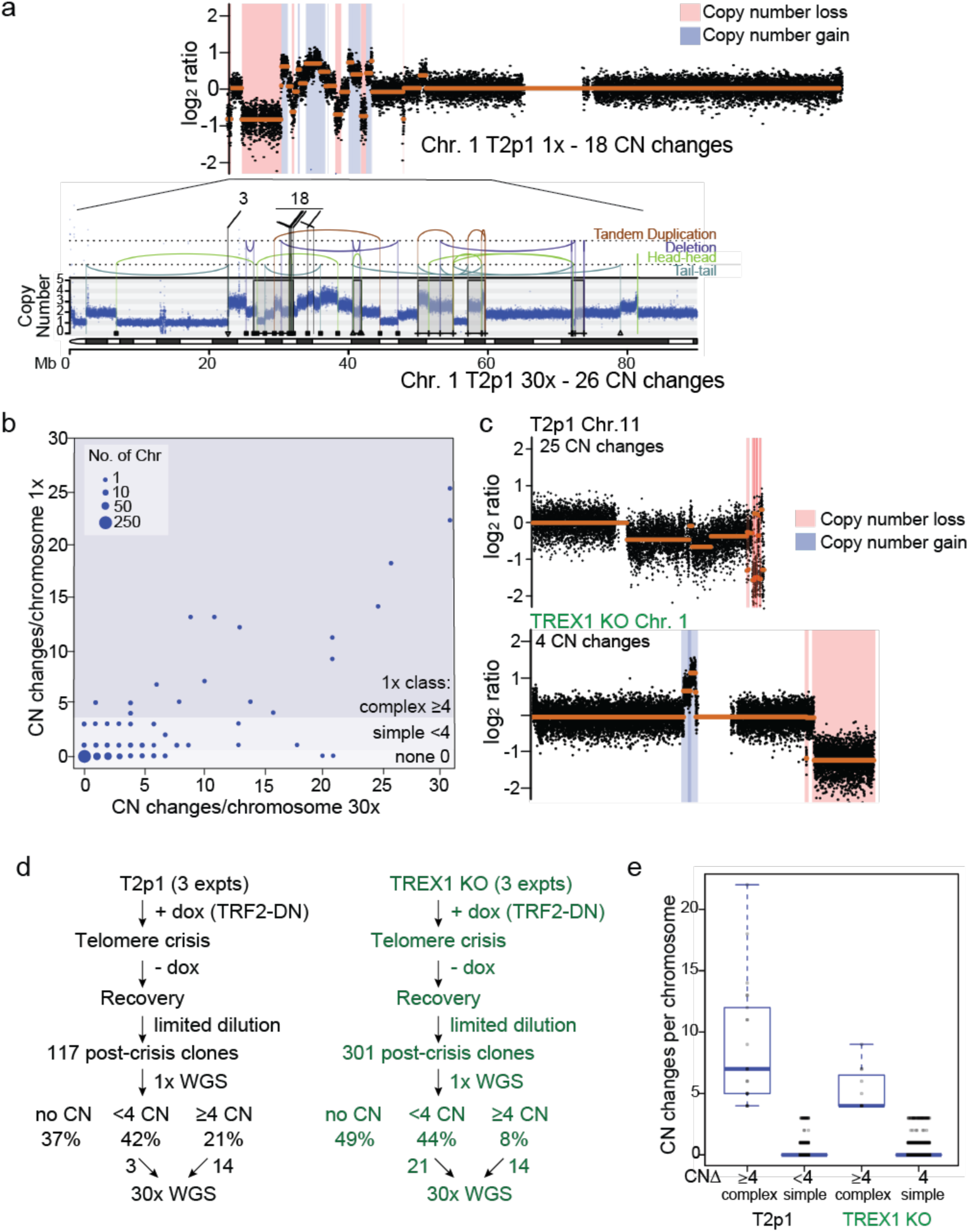
The effect of TREX1 on telomere crisis induced rearrangements. **a**, Example of comparison of 1x and 30x WGS analysis of part of Chromosome 1 in a clone derived from T2p1 cells induced to undergo telomere crisis. Top: QDNAseq analysis of 1x target coverage genomic sequencing data. Regions of CN loss (pink highlight) and CN gain (blue highlight) are indicated. CN profiles are log_2_-transformed. Bottom: DNA CN profile (estimated copy number over genomic windows) and rearrangement joins obtained from Battenberg analysis of 30x target coverage genomic sequencing data. Colored arcs represent joins with the type of rearrangement and orientation as identified on the right. Rearrangement joins are further classified into discrete events as indicated by triangles, squares, and plus symbols (see methods) Interchromosomal rearrangements junctions linking to chromosomes 3 and 18 are indicated. **b**, Comparison of detection of DNA CN changes per chromosome in 1x and 30x WGS of ∼400 chromosomes from post-crisis T2p1 clones. Size of the dots reflect number of chromosomes with the indicated CN changes detected in 1x and 30x WGS. No events, simple events, and complex events are defined on the right. **c,** Examples of DNA copy number profiles (1x (QDNAseq)) of a T2p1 and a TREX1 KO post-crisis clone. **d**, Analysis pipeline and summary of the number of post-crisis T2p1 and TREX1 KO clones isolated from independent telomere crisis experiments, the frequency of simple and complex CN changes detected (1x) and the number of clones selected for 30x WGS. **e,** Box and whisker plot of complex (≥4 CN changes/chromosome) and simple (1-3 CN changes/ chromosome) in rearranged chromosomes from T2p1 and TREX1 KO post-crisis clones. Data derived from 1x (WGS of 117 subclones (2,691 chromosomes) and 301 subclones (6,923 chromosomes) from T2p1 and TREX1 KO clones, respectively. p<0.004 using Student’s t-test.

Comparison of the 1x WGS data obtained from 301 TREX1 KO post-crisis clones with 117 T2p1 clones showed that among clones with CN changes, the frequency of clones with complex events was lower in the TREX1 KO setting (Fig. 1c-e). Furthermore, the average number of CN changes associated with complex events was 2-fold lower in the TREX1 KO setting (Fig. 1c-e; Ext. Data Fig. 1). These results indicate that cells progressing through telomere crisis without TREX1 sustain fewer complex chromosome rearrangements.

Post-crisis clones were screened for copy number changes at 1x and those with a minimum of 4 copy number changes (complex) on at least one chromosome qualified as candidates for sequencing at high coverage (Fig. 1d). Clones with the most changes were sequenced at 30x. In addition, some clones lacking complex events were selected for sequencing at 30x resulting in a total of 17 and 35 clones for T2p1 and TREX1 KO, respectively (Fig. 1d; Ext. Data Fig. 1a). The genomic alterations observed were grouped in four categories: chromothripsis (as defined by^15^; chromothripsis-like which we define here as a chromothripsis pattern with <10 SVs; Breakage-Fusion-Bridge cycles (as defined^16,17^); and a fourth category referred to as Local Jumps. Local Jumps are often unbalanced translocations or large deletions with a locally-derived fragment inserted at the breakpoint^18^ (Fig. 2a).

**Figure 2.**
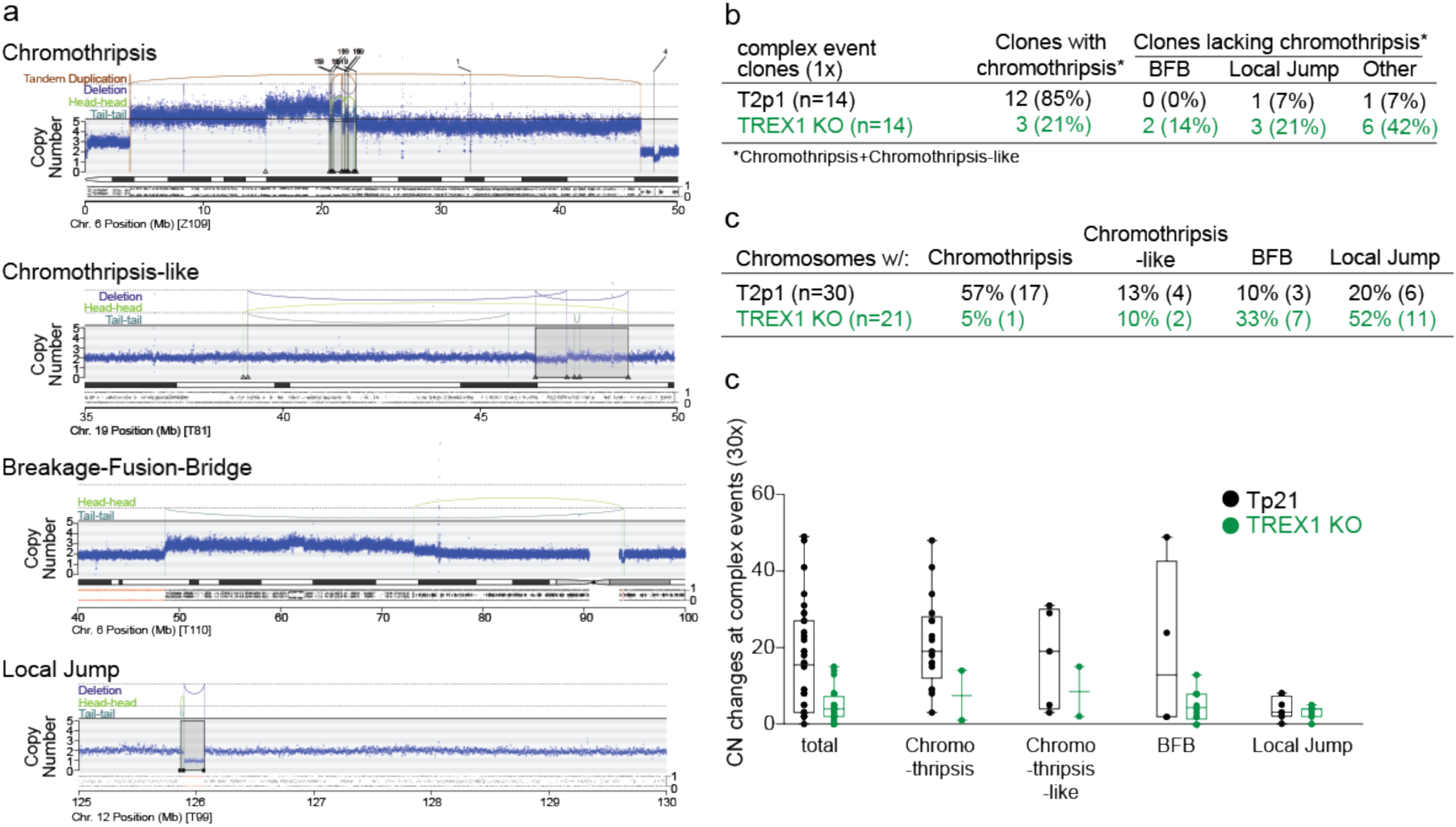
TREX1 promotes chromothripsis. **a**, Examples of chromothripsis, chromothripsis-like, Breakage-Fusion Bridge, and Local Jump patterns in post-crisis clones derived from TREX1 KO cells. DNA CN profiles and rearrangement joins were obtained from 30x target coverage WGS. Annotation as in Fig. 1a. Variant allele frequency (VAF) tracks are shown below the chromosome ideograms. **b,** Summary of number of clones that displayed the types of rearrangements shown in (a) as determined by 30x WGS of 14 T2p1 and TREX1 KO post-crisis clones with complex events observed in 1x WGS. **c,** Summary of the number of chromosomes in post-crisis T2p1 and TREX1 KO clones examined in (b) that display the indicated rearrangements. **d,** Box and whisker plot of the number of CN changes associated with the complex events indicated in post-crisis T2p1 and TREX1 KO clones described in (b).

Of the 14 selected T2p1 post-crisis clones with more than 4 copy number changes in 1x coverage analysis (Fig. 1e; Ext. Data Fig. 1a), 12 (86%) had either chromothripsis or a chromothripsis-like pattern on 30x coverage WGS (Fig. 2a-c). Consistent with telomere dysfunction driven events, chromothripsis was often localized to distal parts of chromosome arms (Ext. Data Fig. 2) In contrast, among the 14 TREX1 KO clones with complex events analyzed by 30x WGS, only three (21%) showed chromothripsis or chromothripsis-like patterns. Taken together with the low-coverage data, these data indicate that chromothripsis is more frequent when cells experience telomere crisis in the presence of TREX1.

The patterns of structural variation in the post-crisis TREX1 KO clones showed that other abnormalities emerge instead of chromothripsis (Fig. 2c). While the majority (57%) of CN changes in the T2p1 clones are classified as chromothripsis or chromothripsis-like, TREX1 KO clones showed enrichment of BFB and Local Jump signatures (Fig. 2c; Ext. Data Fig. 3). Commensurate with this, the number of CN changes per event was lower in the TREX1 KO clones than T2p1 clones (Fig. 2d). The lack of chromothripsis in the TREX1 KO clones was not due to a greater rate of cell death during telomere crisis; in absence of TREX1 apoptosis was observed at the same a low rate as in the T2p1 cells (∼10% showed chromatin changes consistent with apoptosis over a period of 4 days after TRF2-DN induction). The implication of these data is that TREX1 KO cells resolve DNA bridges formed in telomere crisis through simple structural events rather than chromothripsis.

Chromothripsis after telomere crisis is accompanied by kataegis with the hallmark of APOBEC3 editing: clustered and strand-coordinated mutations in cytosine residues in TCA or TCT triplets^4,19^. The ssDNA substrate of APOBEC3 enzymes is formed by TREX1-dependent nucleolytic degradation of the DNA bridges formed in telomere crisis. Based on imaging of Turquoise-tagged RPA70 after TRF2-DN induced telomere fusions (Fig. 3a-c), the ssDNA remnant of resolved DNA bridges appeared either appear to join the primary nucleus or remain outside the nucleus during interphase. In the next mitosis, RPA foci were still detectable and often became incorporated into one of the daughter nuclei. In the majority of cases (47 out of 49 nuclei analyzed), large RPA foci remained detectable for at least 19 h, suggesting that there is a long period in which the ssDNA substrate for APOBEC3 editing persists after DNA bridge resolution.

**Figure 3.**
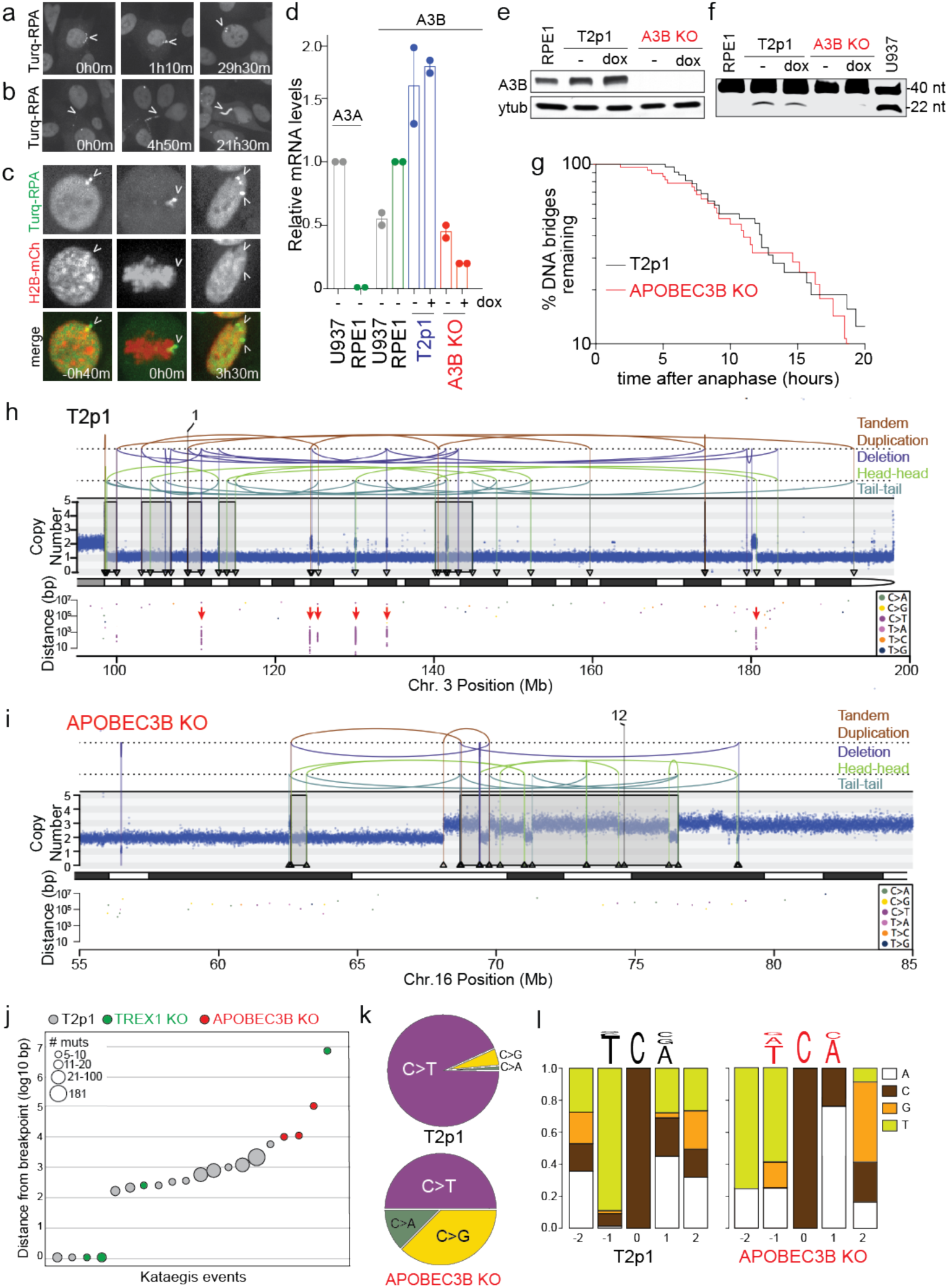
APOBEC3B induces kataegis during telomere crisis. **a**, Stills from live-cell imaging of T2p1 cells expressing mTurquoise2-RPA70 at the indicated time points after doxycycline treatment showing ssDNA joining the primary nucleus. **b,** As in (a) but showing an example of ssDNA remaining outside the nucleus. **c,** As in (a) but showing both H2B-mCherry and representing an example of ssDNA joining a daughter nucleus during mitosis (times before and after mitosis are given). **d,** Relative APOBEC3 mRNA levels in the indicated cell lines based on qRT-PCR. Value of 1 is assigned to U937 cells (APOBEC3A, A3A) or RPE1 cells at baseline (APOBEC3B, A3B). **e,** Immunoblotting for endogenous APOBEC3B before or after 48 hours of dox treatment in the indicated cell lines. **f,** Cytidine deaminase activity assay in the indicated cell lines. Expected DNA fragment sizes are indicated. **g,** Timing of DNA bridge resolution after anaphase in T2p1 and APOBEC3B KO derivative cells expressing H2B-mCherry and mTurquoise2-RPA70 and induced with dox. Data were obtained from >50 DNA bridges per genotype from two independent experiments. **h** and **i**, Examples of DNA CN profile and rearrangement joins of T2p1 (h) and APOBEC3B KO (i) post-telomere crisis clones obtained from 30x WGS. Annotation as in Fig. 1a. Arrows indicated kataegis clusters. **j,** Plot of distance of cytosine mutations to the nearest rearrangement breakpoint. **k,** Distribution of mutation types in the detected kataegis clusters. **l,** Nucleotide context of mutated cytosines. The relevant positions shown are on the pyrimidine (cytosine) strand. The y axis shows the fraction of each nucleotide on the pyrimidine strand. Triplet logos are shown above.

Transcript analysis showed that RPE1 cells express APOBEC3B but not ABOBEC3A (Fig. 3d). The APOBEC3B mRNA levels in T2p1 cells were slightly increased compared to the parental RPE1 cell line but not further induced by telomere damage (Fig. 3d). APOBEC3B was targeted by CRISPR/Cas9 editing (Ext. Data Fig. 4) and loss of APOBEC3B expression was verified by immunoblotting (Fig. 3e). Cytosine deaminase activity in cell extracts became undetectable in APOBEC3B KO cells (Fig. 3f), indicating the APOBEC3B is the major cytosine deaminase in the telomere crisis cell line. The absence of APOBEC3B did not affect the resolution of the DNA bridges formed by dicentric chromosomes (Fig. 3g).

The pipeline of 1x and 30x WGS analysis described above was applied to 375 clones derived from four independent experiments performed with APOBEC3B KO cells (Ext. Data Fig. 5a-c). The percentage of post-crisis clones showing CN changes detectable by 1x WGS and the frequency of clones with either simple or complex events was similar in the absence and presence of APOBEC3B (Ext. Data Fig. 5a). Furthermore, 30x WGS of 23 selected clones showed that the prevalence of chromothripsis and chromothripsis-like events was not affected by the absence of APOBEC3B (Fig. 3h,i; Ext. Data Fig. 5b,c).

As expected, a substantial number of kataegis events involving primarily C to T changes in TCA triplets were observed in the post-crisis wild-type T2p1 clones (Fig. 3j-l). Kataegis was only observed at chromothripsis clusters, and, as expected, all such events were located within 10 kb of the nearest breakpoint and more than half were clusters of more than 10 mutations (ranging from 12-181) (Fig. 3j). Interestingly, kataegis in the T2p1 clones never occurred at the simple BFB and Local Jump breakpoints. Since these simple rearrangements do not require TREX1 (Fig. 2), they may not involve generation of the ssDNA substrate for APOBEC3 editing. In contrast, the APOBEC3B KO post-crisis clones showed only two kataegis events and these events had few mutations (6 and 10 mutations). The spectrum of changes and the nucleotide context of the two kataegis events in the APOBEC3B KO cells appeared different although the number of events was too low to determine whether this distinction is significant (Fig. 3k, l). Collectively, the data provide experimental evidence for the link between APOBEC activity and the generation of Signatures 2 and 13 in the cancer genomes^20^.

The overall frequency of chromothripsis in the APOBEC3B and T2p1 clones was similar and distinct from the lower frequency observed in the TREX1 KO clones (Fig. 4a; Ext. Data Fig. 5b,c). However, chromothripsis events in the APOBEC3B KO clones were distinct from those in the T2p1 clones, showing fewer CN changes per complex event (Fig. 4b,c). The simplest interpretation of this data is that cytosine deamination leads to strand breakage and thereby increases the DNA fragmentation underlying chromothripsis in this setting (Fig. 4e). This strand breakage is not required for DNA bridge resolution since their resolution is not affected by APOBEC3B deletion (Fig. 3g). This lack of effect of APOBEC3B on bridge resolution is consistent with cytosine deamination taking place at a later stage after the bridges have been resolved. It is likely that uracil glycosylases, such as UNG2, and abasic pyrimidine/purine endonucleases, such as APE1^21^ act on the edited ssDNA to result in strand breakage. We propose that in this context, APOBEC3B functions as a cytidine specific initiator of DNA fragmentation. This proposal is consistent with the finding that APOBEC3B overexpression results in DNA damage^22^.

**Figure 4.**
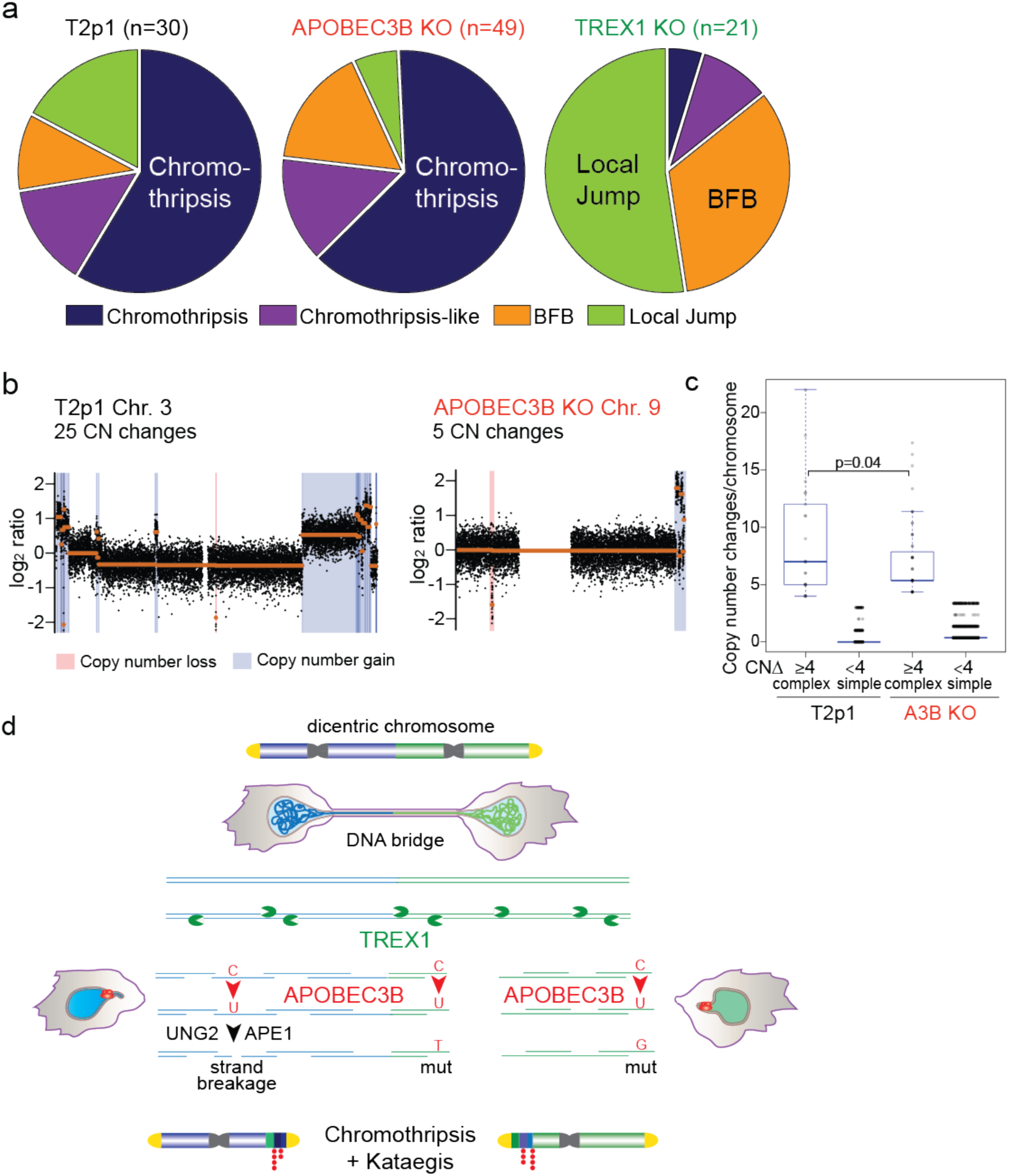
TREX1 and APOBEC3B Determine Genome Instability In Telomere Crisis. **a**, Pie-charts summarizing detected events in the indicated cell lines. Data obtained from 30x WGS. n: Total number of chromosomes with either of the four types of events. **b,** Example of DNA copy number profiles of T2p1 and APOBEC3B KO post-telomere crisis clones from 1x WGS. **c,** Box and whisker plots showing DNA CN changes per altered chromosome in complex and simple events in T2p1 v APOBEC3B KO clones. p=0.04 using Student’s t-test. **d,** Schematic displaying the inferred TREX1- and APOBEC3B-dependent events leading to chromothripsis and kataegis during telomere crisis.

These data established that TREX1 promotes chromothripsis during in vitro telomere crisis. Since TREX1 localizes to DNA bridges, is responsible for the formation of ssDNA, and promotes bridge resolution^4^ we consider it likely that the generation of ssDNA by TREX1 underlies chromothripsis in this system. Furthermore, the finding of APOBEC3 editing at chromothriptic breakpoints in this and other studies^11,23^ is consistent with ssDNA being formed by TREX1 during the process of chromothripsis. We do not know the nature and the frequency of the nicks that provide TREX1 with a starting point for 3’ resection. In addition, it is not yet clear how this 3’ exonuclease leads to generation of ssDNA fragments. One possibility is that ssDNA fragments are released when two TREX1 nucleases meet on opposite strands (Figure 4E). Alternatively, DNA helicases could inadvertently stimulate the release of ssDNA fragments or ssDNA could undergo breakage due to physical stress. We also do not know how the ssDNA fragments are converted into the ds DNA fragments that eventually are combined into the chromothripsis region. Our findings imply that the most spectacular genome alteration in cancer is due to the inappropriate action of two enzymes that evolved to defend mammalian genomes against viral invasion.

## Methods

### Data reporting

No statistical methods were used to predetermine sample size. The experiments were not randomized and the investigators were not blinded to allocation during experiments and outcome assessment.

### Cell Culture Procedures and Plasmids

RPE1-hTERT and U937 cells were obtained from the American Type Culture Collection (ATCC). RPE1-hTERT cells were cultured in a 1:1 mixture of Dulbecco’s Modified Eagle Medium (DMEM) and Ham’s F-12 medium (Gibco) (DMEM/F12). Phoenix virus packaging cells were grown in DMEM. U937 cells were grown in RPMI-1640 medium. All media were supplemented with 10% fetal bovine serum (Gibco), 100 U/ml penicillin/streptomycin (Life Technologies), and 2.5 mM L-glutamine (Life Technologies). T2p1 cells and its TREX1 KO derivatives were described previously^4^. Doxycycline was used at 1 µg/ml.

Target sequence for CRISPR/Cas9 mediated gene knockouts identified by ZiFit (http://zifit.partners.org): sgAPOBEC3B #1: GGAGCCCTCCCGGGAGTCCT; sgAPOBEC3B #2: GGGGTGCTCAGACAGGAATT. Plasmids containing sgRNAs (Addgene 41824) and a human codon-optimized Cas9 (Addgene 41815) were co-nucleofected into target cells by nucleofection (Lonza apparatus). 700,000 cells were mixed with electroporation buffer (freshly mixed 125 mM Na_2_HPO_4_, 12.5 mM KCl, 55 mM MgCl2 pH 7.75), 5 µg Cas9 plasmid, and 5 µg gRNA plasmid, transferred to an electroporation cuvette (BTX), and electroporated with program T23 for T2p1 cells. Cells were then allowed to recover for 48 h before a second round of electroporation. Successful CRISPR/Cas9 editing was confirmed at the polyclonal stage by mutation detection with the SURVEYOR nuclease assay (Transgenomic). The regions surrounding the Cas9 cut sites were PCR amplified (JM661: GCCTCAGAATTCAGTTCTACCAT, JM662: TCGATTGAAGTTTGGGTGGT; JM657: AAGGAGCTTCAATGGCAAGA, JM658: GCATGAATTGCTGACCTTCA), melted, and reannealed. Reannealed PCR products were incubated with the SURVEYOR nuclease for one hour at 42°C and analyzed on a 2% agarose gel with ethidium bromide. Clones were isolated by limiting dilution and screened for APOBEC3B deletion by PCR. Inversions resulting from successful sgA3B #1 and sgA3B #2 cutting were identified using primers JM662 and JM680: CATAGTCCATGATCGTCACGC. Deletion of the wt APOBEC3B allele was confirmed using primers JM679: TGTATTTCAAGCCTCAGTACCACG and JM680. Biallelic targeting was verified by Western blotting and sequencing of TOPO-cloned PCR products.

### Immunoblotting

For immunoblotting, cells were harvested by trypsinization and lysed in 1x Laemmli buffer (50 mM Tris, 10% glycerol, 2% SDS, 0.01% bromophenol blue, 2.5% β-mercaptoethanol) at 10^7^ cells/ml. Lysates were denatured at 100°C and DNA was sheared with a 28 1/2 gauge insulin needle. Lysate equivalent to 10^5^ cells was resolved on 8% or 10% SDS/PAGE (Life Technologies) and transferred to nitrocellulose membranes. Membranes were blocked in 5% milk in TBS with 0.1% Tween-20 (TBS-T) and incubated with primary antibody overnight at 4°C, washed 3 times in TBS-T, and incubated for 1 h at room temperature with horseradish-peroxidase-conjugated sheep anti-mouse or donkey anti-rabbit secondary antibodies. After three washes in TBS-T, membranes were rinsed in TBS and proteins were developed using enhanced chemiluminescence (Amersham).

The following primary antibodies were used: anti-APOBEC3B (rabbit monoclonal, Abcam, ab184990) and anti-γ-tubulin (mouse monoclonal, Abcam, ab11316).

### Live-cell Imaging and quantitation

Live-cell imaging of mCherry-H2B marked cells was performed as described previously^4^. Chromatin bridge resolution was determined by manually tracking pairs of daughter cells. Bridge resolution was inferred to take place when the base of the bridge became slack and/or recoiled. RPA and APOBEC3B were tracked based on mTurquoise2-RPA70 and APOBEC3B-mTurquoise2.

### Quantitative PCR

Random hexameric primers, avian myeloblastosis virus reverse transcriptase (AMV RT; Roche) were used to synthesize cDNA from total RNA (2.5 µg) template. cDNA levels were quantified by PCR using a Roche Lightcycler 480 instrument as described^24^. In brief, reactions were performed in 384-well plates with each well containing 7.5 µl 2x probe master mix (Roche), 1.25 µl H2), 1.05 µl primers (5 µm each), 0.2 µl UPL probe (Roche) and 5 µl cDNA. Reactions were incubated at 95°C for 10 min, then 40 cycles of 95°C for 10 s, 58°C for 15 s, then 72°C for 2s. APOBEC3A qPCR was performed using primer 1: GAGAAGGGACAAGCACATGG and primer 2: TGGATCCATCAAGTGTCTGG. APOBEC3B qPCR was performed using primer 1: GACCCTTTGGTCCTTCGAC and primer 2: GCACAGCCCCAGGAGAAG. Ct values were calculated using the Lightcycler 480 software. cDNA was synthesized and qPCR was performed in triplicate for each sample.

### In vitro deamination assay

Cells were lysed in 25 mM HEPES, 5 mM EDTA, 10% glycerol, 1 mM DTT, and protease inhibitor. Protein concentrations were equalized by cell counting prior to lysis. Deamination reactions were performed at 37° C using the APOBEC3B probe (5’ IRDYE800-ATTATTATTATTATTATTATTTCATTTATTTATTTATTTA 3’) in a 10x UDG reaction buffer consisting of 1.25 µl RNaseA (0.125 mg/ml), 1 µl probe (0.2 pmol/µl), 16.5 µl cleared lysate and uracil DNA glycosylase (UDG; NEB, 1.25 units). Abasic site cleavage was induced by the addition of 100 mM NaOH and incubation at 95° C. Reaction products were migrated on 15% urea-TBE gels and imaged on an Odyssey CLx Imaging System (Licor).

### X-ten Sequencing and Mapping

Genomic DNA sequencing libraries were synthesized on robots and cluster generation and sequencing were performed using the manufacturer pipelines. Average sequence coverage across the samples was 37.3x (range, 23.5 – 47.8x). Sequencing reads were aligned to the NCBI build 37 human genome using the BWA mem algorithm (version 0.7.15;^25^) to create a BAM file with Smith-Waterman correction with PCR duplicates removed [http://broadinstitute.github.io/picard/].

### Mutation Calling

Point mutations were called using CaVEMan version 1.11.2^26^ with RPE-1 as reference. A simple tandem repeat filter was applied first to remove variants observed less than five times or were seen in less than 10% of the reads. Also, a variant was considered only if observed in both forward and reverse strands. To enrich for high-confidence somatic variants, variants were further filtered by removing: known constitutional polymorphisms using human variation databases: Ensembl GRCh37, 1000 genomes release 2.2.2, ESP6500 and ExAC 0.3.1.

Raw mutations were filtered using a homopolymer filter. Mutations which had a homopolymer repeat of at least six bases on either side of the mutation and where the mutated base was same as the base of the homopolymer repeat(s) were removed. A soft-clip filter was used in a similar way, mutations where more than half of the supporting reads were softclipped were removed.

### Copy number analysis

We detected DNA copy number aberrations by shallow WGS at 1x (average 1.3x) using QDNAseq^13^. The genome was divided into bins of 15kb and the method used for the callBins was “cutoffs” for deletion = 0.5, loss = 1.2, gain = 2.5, amplification = 10. A blacklist of copy number changes repeated in the same regions in at least 10% of the samples was reported and removed from the final copy number data at 1x. We counted copy number changes per chromosome for each sample and selected those with more than four copy number changes to sequence at 30x. Samples with more than four copy number changes were likely to have real events which was confirmed by X ten sequencing at 30x.

We used both Ascat^27^ and Battenberg (https://github.com/cancerit/cgpBattenberg) to extract copy number data from 30x WGS. Ascat was used assuming ploidy of 4 for subclonal event identification and to overall enhance aberrations for easier data manipulation. Battenberg analysis was performed using ploidy of 2 which was consistent with the QDNAseq settings for direct comparison of the data from the two algorithms.

### Event identification

Events were defined through regions with high density rearrangement breakpoints. A minimum of 4 breakpoints spaced 2Mb apart was identified as an event. The rest of the rules applied for the identification of the events were related to the propagation of the rearrangements. When one breakpoint of a rearrangement was part of an event while the second wasn’t because of the distance rules (above) applied, the two breakpoints were merged into the same event. When the breakpoints of the same rearrangement belonged to different events, they were merged into one event. To graphically distinguish between different events on our plots we annotated breakpoints of events using different shapes at the bottom tips of their breakpoints (Fig. 3e,f).

### Rearrangement Calling and Chromothripsis

To call rearrangements we applied the BRASS (breakpoint via assembly) algorithm, which identifies rearrangements by grouping discordant read pairs that point to the same breakpoint event (github.com/cancerit/BRASS). Post-processing filters were applied to the output to improve specificity (blacklisted recurrent breakpoints in 10% of samples). Complex chromothripsis clusters were called according to the criteria from^26^. 1. A minimum of 4 breakpoints spaced 2Mb apart was considered an event of high density. 2. Oscillating copy number stages were mostly detected but non-conventional chromothripsis was also seen. 3. Multiple chromosomes retained loss of heterozygosity across chromosomes. 4. 1x WGS data analysis confirms prevalence of rearrangements. 5. The type of fragment joins in chromothripsis should be uniformly distributed. However, the chromothripsis events involve fairly low numbers of intra-chromosomal rearrangements, which would decrease power in a uniform multinomial distribution. 6. Ability to walk the derivative chromosome was not an applicable rule, as chromothripsis takes place on chromosomes with preceding duplication through BFBs^28^.

Another category of events identified during this study was the Chromothripsis-like events, defined here as having <10 SVs but patterns consistent with chromothripsis. Local jumps seen mainly in TREX1 KO clones are defined according to a prior report^18^. They consist of rearrangements reaching into one or more distant regions of the genome. Simple examples of these events include unbalanced translocation or large deletion with a locally-derived fragment inserted at the breakpoint, but there was also an extensive range of more complex patterns (local n jumps)

### Kataegis

Kataegis mutation clusters were detected according to^29^. Similar to the identification of events, mutations spaced ≤2 kb apart were treated as a single mutagenic event. Groups of closely spaced mutations (at least four mutations) were identified, such that any pair of adjacent mutations within each group was separated by less than 2 kb. To identify clusters that were unlikely to have formed by the random distribution of mutations within a genome, we computed a *P* value for each group. Each group with *P* ≤ 1. 10^−4^ was considered a bona fide mutation cluster. A recursive approach was applied, i.e., all clusters passing *P*-value filtering were identified, even if a cluster represented a subset within a larger group that did not pass the *P*-value filter.

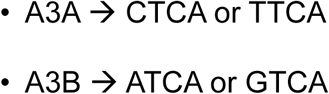

TCA enrichment was calculated and significance was assessed using Fisher’s exact test

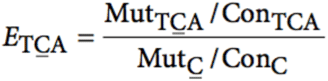

The enrichment of YTCA and RTCA was calculated and significance was assessed using chi-squared test based on the expected YTCA and RTCA.

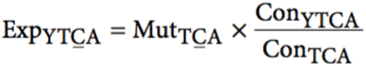

Whhere ConTCA = TCA occurrences

Enrichment of C->G and C->T mutations in the TCA context compared to other contexts and normalized it by how many times the motif occurs in the genome.

### Statistical model for kataegis association with genotype

We found a statistically significant relationship when comparing APOBEC KO to T2p1 kataegis clusters by applying the negative binomial distribution to test how kataegis clusters are related to rearrangements across genotypes. Our model showed that APOBEC_KO samples contain a high enough number of breakpoints expected to see in kataegis clusters. The same is not true for TREX1 KO samples. Model examined is:

kat$gen_factor <- factor(kat&genotype, levels = c(“crisis-control”, “APOBEC_KO”, “TREX1_KO”))

summary(glm.nb(kat_clusters ∼ gen_factor + offset(log(SV)), data = kat))

## ACCESSION NUMBERS

The accession number for the genome sequence data reported in this paper is European Nucleotide Archive (http://www.ebi.ac.uk/ena/, hosted by the EBI): Study: PRJEB23723, Secondary accession ERP105494 (https://www.ebi.ac.uk/ena/data/view/PRJEB23723).

## Acknowledgements

We thank Sally Dewhurst for insightful comments on this manuscript. Research reported in this publication was supported by grants from the National Cancer Institute (R35CA210036) and from the Breast Cancer Research Foundation to T.d.L.. T.d.L. is an American Cancer Society Rose Zarucki Trust Research Professor. J.M. is supported by grants from the National Cancer Institute (R00CA212290), an MSK Cancer Center Support Grant/Core Grant (P30 CA008748), and the V Foundation for Cancer Research.

## Author contribution

J.M., T.d.L., and P.J.C. conceived and designed the study; J.M., A.C., and A.D. performed the experiments and generated and analyzed the data. J.M., A.C., and T.d.L. wrote the manuscript with contributions from P.J.C. All authors approved the final manuscript.

---

**Ext. Data Fig. 1.**
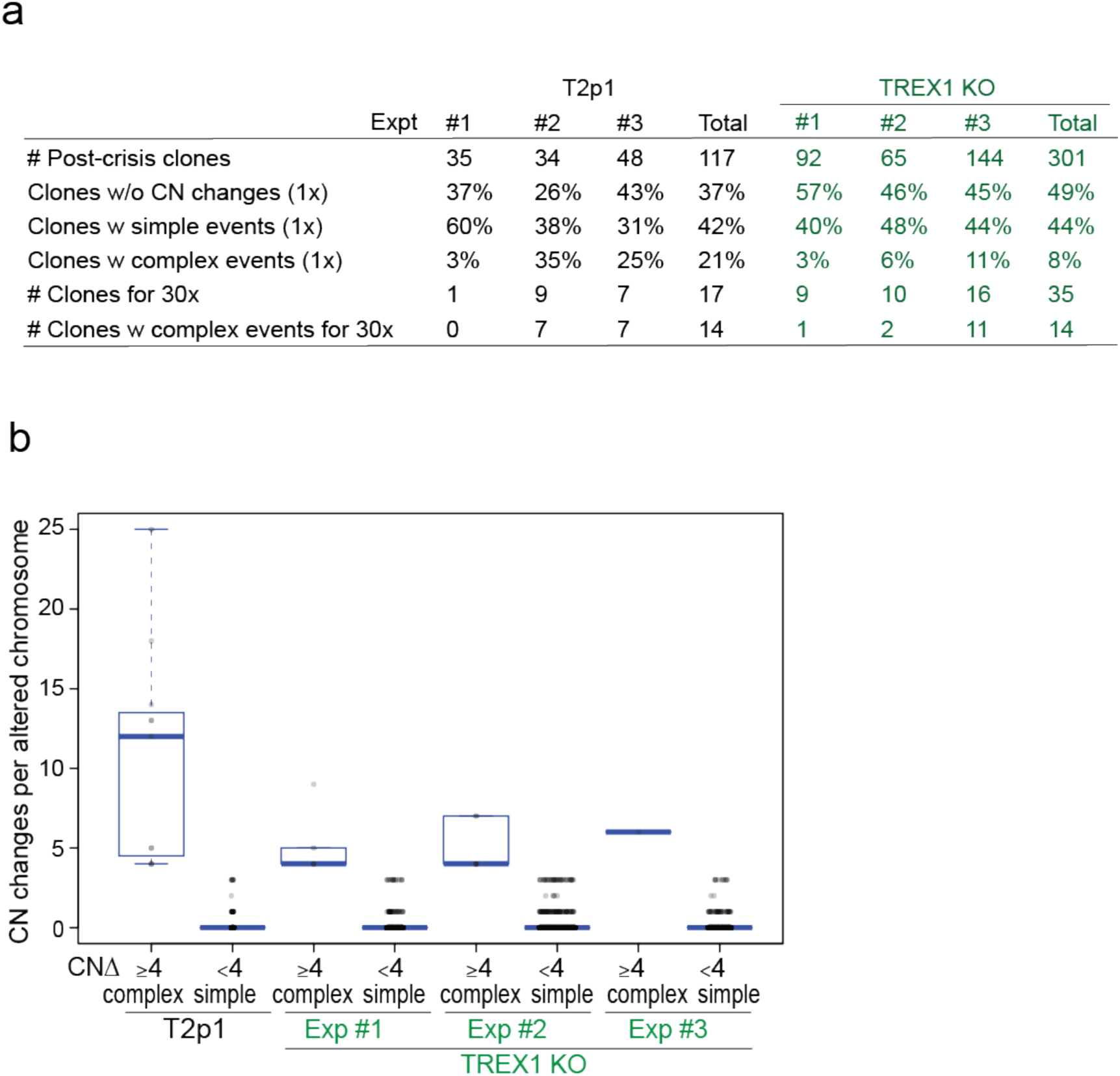
TREX1 affects CN alterations detected by 1x WGS analysis. **a**, Summary of the number of post-crisis T2p1 and TREX1 KO clones isolated from independent telomere crisis experiments, the frequency of simple and complex CN changes detected (1x) and the number of clones selected for 30x WGS. **b,** Box and whisker plot of complex (≥4) and simple (1-3) CN changes in rearranged chromosomes from T2p1 and TREX1 KO post-crisis clones. Data derived as in Fig. 1f.

**Ext. Data Fig. 2.**
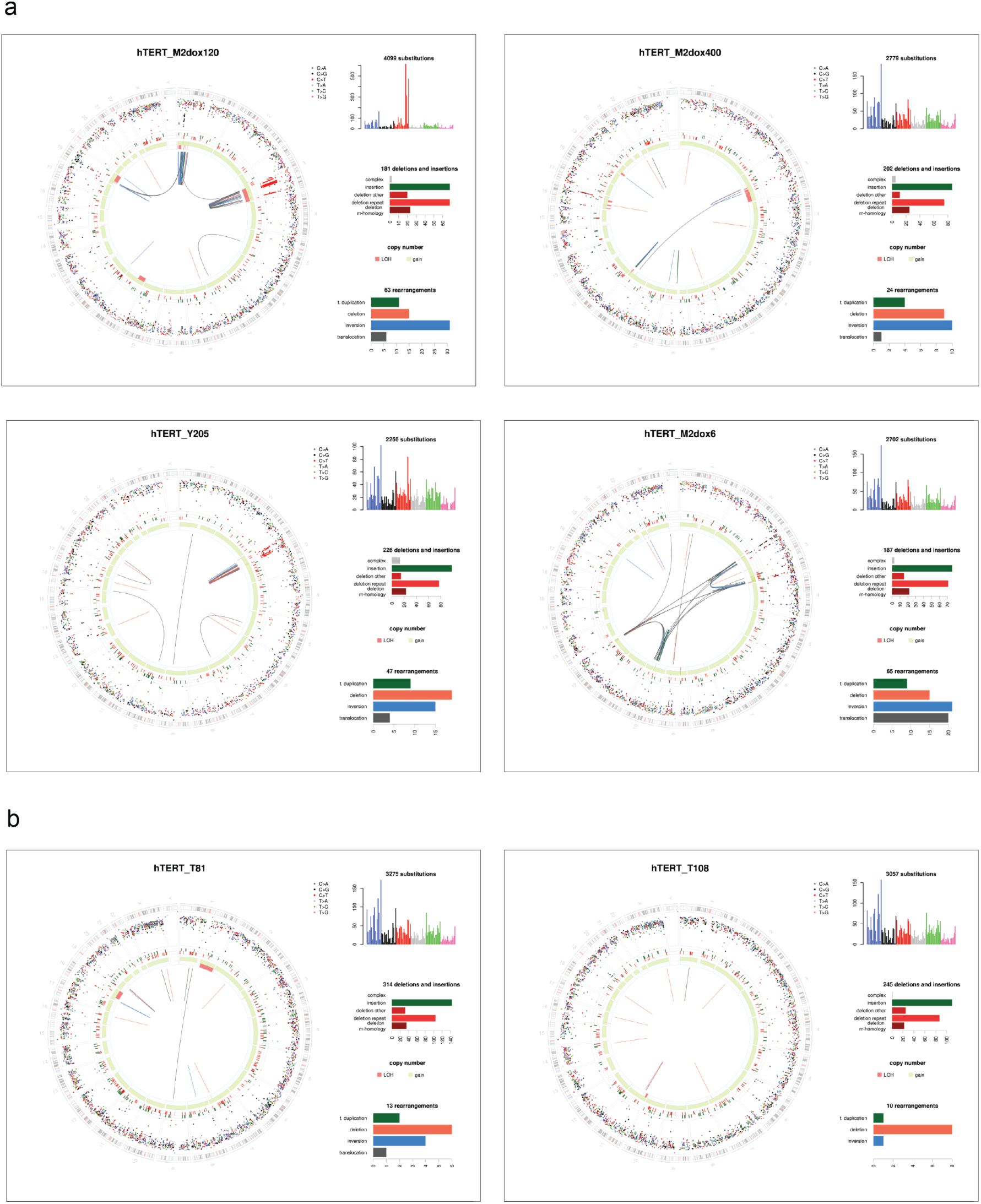
Circos plots of genome alterations in T2p1 post-crisis clones. **a**, Four T2p1 post-crisis clones with complex events identified by 1x WGS were analyzed by 30x WGS and their associated genome plots are shown. Circos plots show somatic mutations including substitutions (outermost, dots represent 6 mutation types: C>A, blue; C>G, black; C>T, red; T>A, grey; T>C, green; T>G, pink), indels (the second outer circle, color bars represent five types of indels: complex, grey; insertion, green; deletion other, red; repeat-mediated deletion, light red; microhomology-mediated deletion, dark red) and rearrangements (innermost, lines representing different types of rearrangements: tandem duplications, green; deletions, orange; inversions, blue; translocations: grey). The number of detected base substitutions, indels, and rearrangements are shown to the right of each panel. **b,** Genomic information on two post-crisis TREX1 KO clones displayed as in (a).

**Ext. Data Fig. 3.**
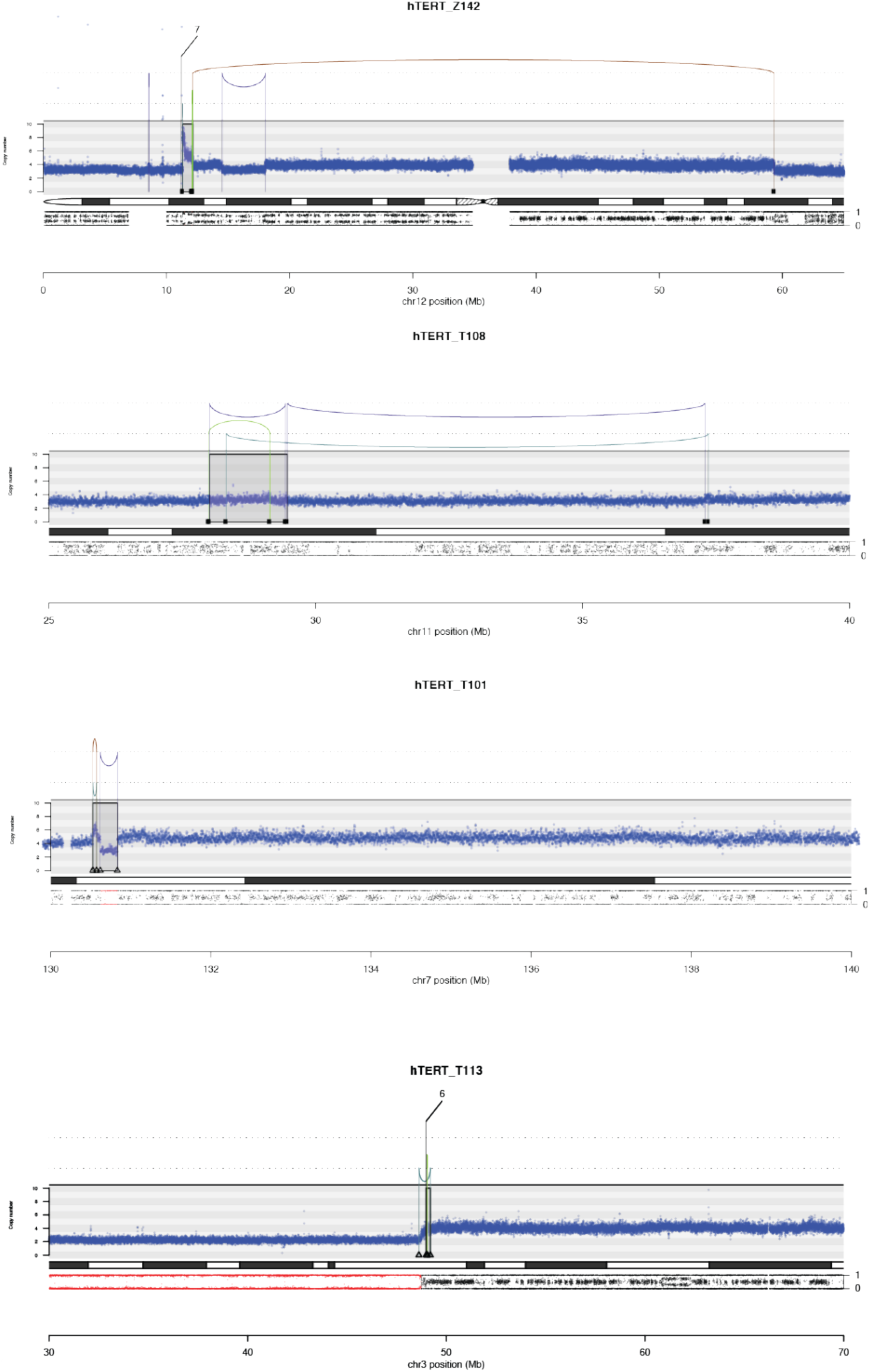
Chromothripsis-like, BFB and Local Jump events in TREX1 KO post-crisis clones. Four TREX1 KO post-crisis subclones with complex (Z142, T113) events, simple (T108) events or no rearrangements (T101) identified by 1x WGS were analyzed by 30x WGS. DNA CN profiles and rearrangement joins were obtained from Battenberg analysis of 30x target coverage genomic sequencing data. Annotation as in Fig. 1a. Variant allele frequency (VAF) tracks are shown below the chromosome ideograms. Examples show a chromothripsis-like event in Z142, a local n jump in T108, a local 3 jump in T101, and a BFB event in T113.

**Ext. Data Fig. 4.**
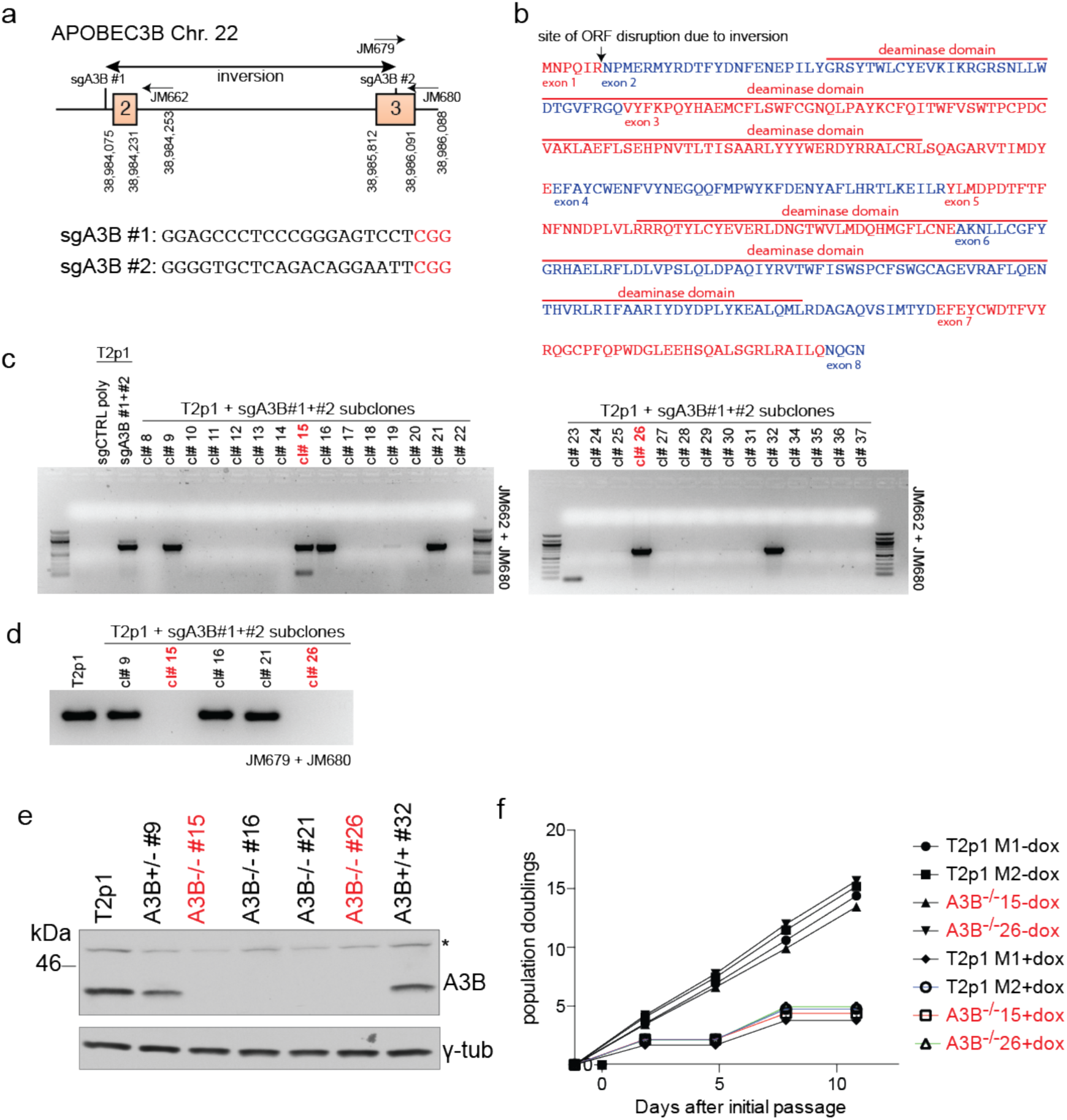
CRISPR/Cas9 Gene Editing of APOBEC3B. **a**, Schematic of the APOBEC3B locus showing landmarks relevant to CRISPR editing. sgRNA sequences used for CRISPR editing are shown below. Protospacer adjacent motifs are marked in red. **b,** APOBEC3B amino acid sequence showing exon boundaries, catalytic domains, and predicted gene disruption from CRISPR editing. **c,** PCR screening identifies clones harboring at least one copy of a CRISPR-generated inversion in the APOBEC3B locus. Clones used for subsequent experiments are marked in red. **d,** PCR screening confirms biallelic disruption of the endogenous APOBEC3B locus. Clones used for subsequent experiments are marked in red. **e,** Immunoblot for APOBEC3B and γ-tubulin shows absence of APOBEC3B in 4 CRISPR-edited clones. Clones #15 and #26 were selected for further this study. Asterisk marks a cross-reacting polypeptide. **f,** Proliferation of the APOBEC3B CRISPR KO clones with and without doxycycline induction of telomere fusions.

**Ext. Data Fig. 5.**
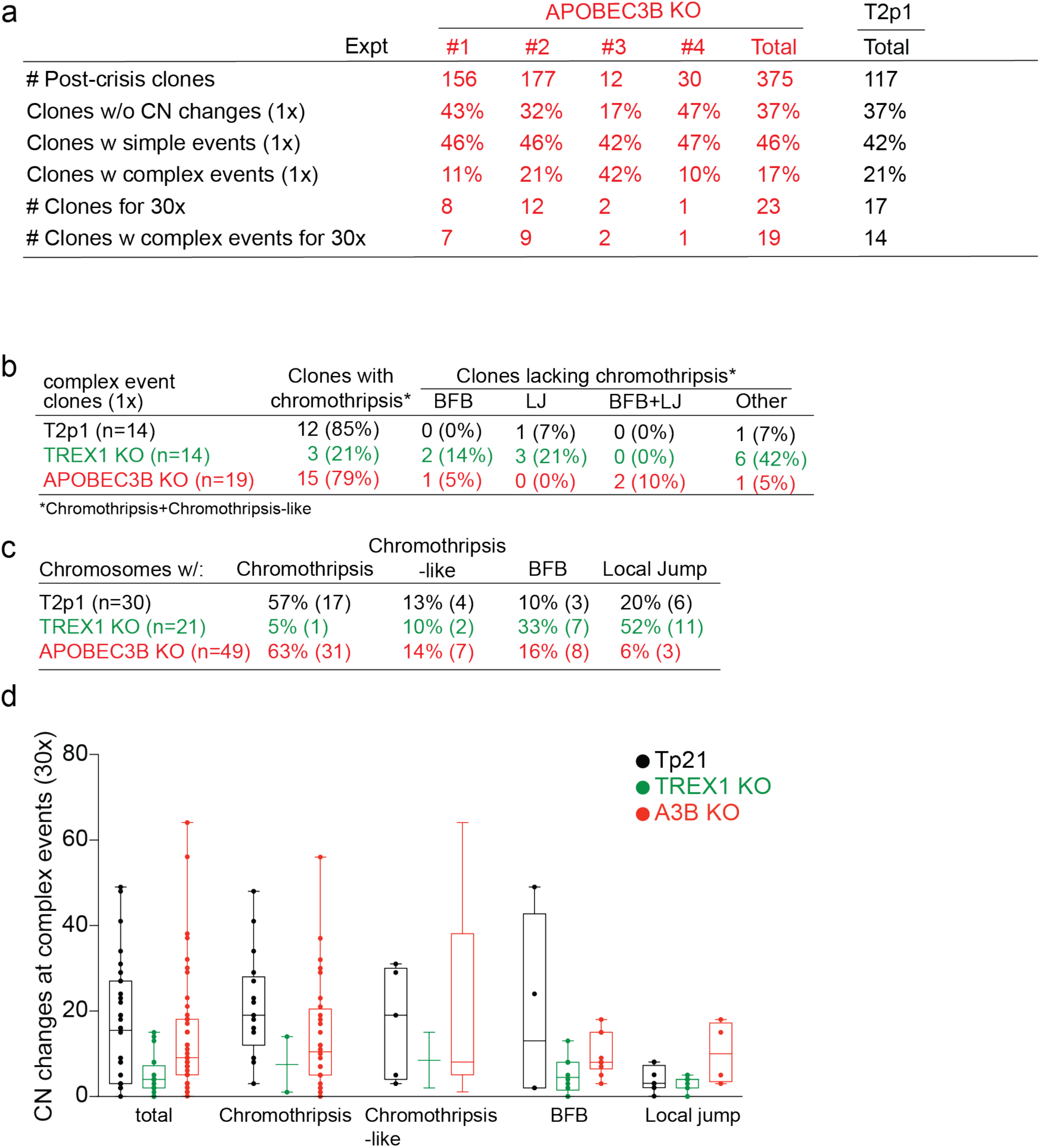
1x and 30x WGS information on APOBEC3B KO post-crisis clones. **a**, Summary of the number of APOBEC3B KO clones isolated from independent telomere crisis experiments, the frequency of simple and complex CN changes detected (1x) and the number of clones selected for 30x WGS. Parallel information on the T2p1 (Fig. 1e) is provided for comparison. **b,** and **c,** Information of complex events in APOBEC3B clones as in Fig. 2b and Fig. 2c. Parallel information on T2p1 and TREX1 KO post crisis clones is provided (from Figs. 1 and 2). **d,** Box and whisker plot displaying the number of CN changes associated with the complex events indicated in APOBEC3B KO clones together with T2p1 and TREX1 KO information (from Fig. 2c).

